# Lack of tolerance development following sublethal Cry1 proteins exposure in *Spodoptera exigua^1^*

**DOI:** 10.1101/2025.10.29.685268

**Authors:** Sandy Nicole Valdiviezo-Orellana, Baltasar Escriche, Patricia Hernández-Mártinez

## Abstract

**BACKGROUND:** The insecticidal proteins derived from *Bacillus thuringiensis* Berliner (Bt) have been effectively employed in controlling lepidopteran pests, notably in transgenic crops targeting *Spodoptera* species. However, concerns have arisen regarding the long-term efficacy due to the emergence of tolerant and resistant insect populations. Prior research suggested that repeated exposures to Bt may contribute tolerance, but the specific effects of sequential exposure to purified Cry1 proteins remain unclear. This study aimed to assess whether prior exposure of *Spodoptera exigua* (Hüber, 1808) neonate larvae to sublethal concentrations of Cry1Ab and Cry1Ca proteins would heighten their tolerance upon subsequent exposure, and whether such effects would extend to their offspring.

**RESULTS:** Pre-exposure to Cry1Ab did not affect larval responses to the toxin. For Cry1Ca, a slight increase was observed under one treatment condition, but the effect was not considered biologically relevant in practical terms. Similarly, transgenerational analysis revealed no enhancement of tolerance; rather, there was a negative impact on the offspring’s response in some cases.

**CONCLUSION:** These findings indicate that although previous studies have documented that sublethal contact with bacterial preparations may significantly affect the insect tolerance, exposure to purified Cry proteins is unlikely to lead to the development of tolerance in *S. exigua*. Therefore, our findings suggest that sublethal exposure to these Cry1 proteins may not significantly affect the long-term efficacy of Bt-based pest management strategies relaying on them.

## 1 INTRODUCTION

*Spodoptera exigua* (Hüber, 1808), commonly known as the beet armyworm, is a polyphagous pest of global distribution that affects a wide variety of crops, including vegetable, field and flowers. The insect can feed on over 170 plant species, principally consuming leaves and fruits, causing significant damage and substantial economic losses.^1,2^ Among the strategies employed for its management, products derived from *Bacillus thuringiensis* (Bt) (Berliner,1915) have proven to be effective. These include transgenic plants expressing Cry and Vip proteins, encoded by genes present in Bt, to control *Spodoptera* spp.^3–5^ By 2020 the global area of transgenic crops reached approximately 186 million hectares, mainly consisting of cotton, maize, soybeans, and canola.^6^ Although transgenic crops have significantly benefited farmers by increasing production and profits, their extensive and, sometimes, indiscriminate use has provided selective pressure on pest populations, leading to the development of resistances. Until 2023, 26 cases of field-evolved practical resistance and 17 cases of early warning of resistance to Bt crops have been reported.^7–28^ Such cases have been primarily attributed to modifications in target receptors,^29^ although it has also been less frequently linked to other mechanisms such as toxin sequestration, alterations in proteolytic processing, and more efficient repair of damaged midgut epithelial cells.^30,31^ In addition, the potential involvement of the immune system in the development of resistance to Bt has been examined. While the evidence remains inconclusive, elevated immune activity appears to play an important role in enhancing survival rates against Bt proteins.^32^ It has been suggested that when an insect is exposed to sublethal doses of Bt vegetative cells, spores and endotoxins or heat-killed bacteria cells it can mount a stronger immune response upon subsequent exposure. This enhanced response, known as immune priming, can lead to increased survival or tolerance to subsequent lethal treatments. Evidence of immune priming has been observed in species such as *Bombyx mori*,^33^ *Ephestia kuehniella*,^34^ *Galleria mellonella*,^35^ *Rhynchophorus ferrugineus*,^36^ *Tribolium castaneum*,^37,38^ and *Tenebrio molitor*.^39^ This increased tolerance has been associated with changes in the regulation of specific immune components, such as the Toll and IMD pathways, hemocyte proliferation, enhanced phagocytic activity, and higher expression of antimicrobial peptides.^34–36,38,40^ Furthermore, this decreased on susceptibility to Bt can be inherited by offspring, a phenomenon known as transgenerational immune priming (TgIP). TgIP has been reported in *T. castaneum*,^41,42^ *T. molitor*,^43–45^ and *Trichoplusia ni*.^46^

In the case of *S. exigua*, knowledge remains limited. Recent studies have stated that priming with heat-killed *Escherichia coli* via hemocoelic injection enhances survival upon subsequent challenges with lethal doses of *Acidovorax citrulli*, *Beauveria bassiana*, and *Xenorhabdus hominickii*. These findings suggest that prior exposure to non-pathogenic bacteria can indeed elicit an immune-priming effect in *S. exigua*.^47^ While immune priming in *S. exigua* has been documented, its relevance to *B. thuringiensis* and its insecticidal proteins remains unclear. Moreover, most existing studies involve hemocoelic inoculation, leaving the effects of oral exposure largely unexplored in this context. Previous research has demonstrated that sublethal exposure to isolated Bt proteins, such as Cry and Vip, in *S. exigua* larvae induces changes in the expression of immune-related genes.^48–50^ However, whether these responses can trigger similar priming effects, altering the tolerance to Bt proteins and potentially undermining the effectiveness of Bt-based pest control strategies needs further clarification. Then, this study aimed to assess whether prior exposure of neonate *S. exigua* larvae to purified Cry1Ab and Cry1Ca toxins would heighten their toxicological response upon subsequent exposure and whether such effects would extend to their offspring.

## 2 MATERIALS AND METHODS

### 2.1 Insects

All the experiments were conducted using a laboratory strain of *S. exigua* reared on an artificial diet,^51^ under controlled conditions of 16/8 h light/dark at 25 ± 3 °C and 70 ± 5% of relative humidity. This strain has been maintained for a minimum of 10 years without exposure to *B. thuringiensis* insecticidal proteins.

### 2.2 Expression and purification of Cry proteins

Cry1Ab and Cry1Ca proteins were obtained from a recombinant *Escherichia coli* strain kindly supplied by R. de Maagd (Cry1Ab plasmid pBD140, Cry1Ca plasmid pBD150). Protein production, inclusion-bodies purification, solubilization, and protoxin activation by trypsin were performed according to the protocol described by Sayyed et al.^52^ For Cry1Ca protein, an additional purification step was necessary. The protein was purified by anion-exchange chromatography utilizing the ÅKTA Explorer 100 System, as described previously by Crava et al.^48^ Fractions were collected and screened for the presence of Cry1Ca using 12% sodium dodecyl sulfate polyacrylamide gel electrophoresis (SDS-PAGE). The activated proteins were analyzed by 12% SDS–PAGE and were subsequently stored at −20°C until further use. The quantification of Cry1Ab and Cry1Ca used in bioassays was performed via densitometry following SDS-PAGE electrophoresis, using BSA as a standard and TotalLab 1D (v13.01) software.

### 2.3 Intragenerational bioassays

The impact of prior exposures to sublethal concentrations of Cry1 proteins on subsequent challenges was analyzed by determining the percentage of growth inhibition. First-instar larvae were initially exposed to several sublethal concentrations of Cry proteins using the surface contamination method.^53^ Larvae were maintained on this treated diet for five days. Mortality and weight of surviving larvae were recorded, before being transferred to non-treated diet until fourth instar, at which point they were exposed to the Cry proteins again. For the challenge, larvae were exposed to different protein concentrations for 24 hours using the same surface contamination methodology. Subsequently, their weights were recorded to calculate the percentage of growth inhibition (%GI) following the methodology described by Herrero et al.^54^ Additionally, a control group was included, wherein larvae were exposed only to the solubilization buffer. Three independent biological replicates were performed for each protein.

For the Cry1Ab protein, two different concentrations, 7.5 ng/cm² and 75 ng/cm², were used for the first exposure, and three different concentrations, 7.5 ng/cm², 75 ng/cm² and 750 ng/cm², for second exposures, based on the IC_50_ values reported by Hernández-Martínez et al.^55^ For the Cry1Ca protein, neonates were initially exposed to 1 ng/cm² and 10 ng/cm², also considering the IC₅₀ values. Subsequently, the second exposure was performed using three different concentrations, 10 ng/cm², 100 ng/cm², and 1000 ng/cm², to evaluate the effect on growth inhibition.

### 2.4 Transgenerational bioassays

To evaluate whether there was an enhancement in toxicological response among larvae originating from a pre-exposure generation, weight in neonates (L1) offspring and percentage of growth inhibition in L4 offspring were measured. Following the methodology as previously described, neonate larvae were exposed to sublethal concentrations of the protein for five days. Then, larvae were transferred to non-treated diet and allowed to produce the F1 generation. The offspring were then divided into two groups: neonate larvae designated for the challenge and the larvae to be maintained until reaching the L4 instar, where another challenge would be performed. The neonate larvae in the first group were exposed to different concentrations of the Cry proteins for 5 days. After exposure, they were weighed and compared with untreated larvae. The second group of larvae, upon reaching the L4, were treated with different concentrations of the protein for 24 hours, and growth inhibition was calculated. Three independent biological replicates were performed for each protein to ensure reliability and consistency in the results. A schematic overview of the methodology described above is presented in Figure 1.

**Fig. 1.**
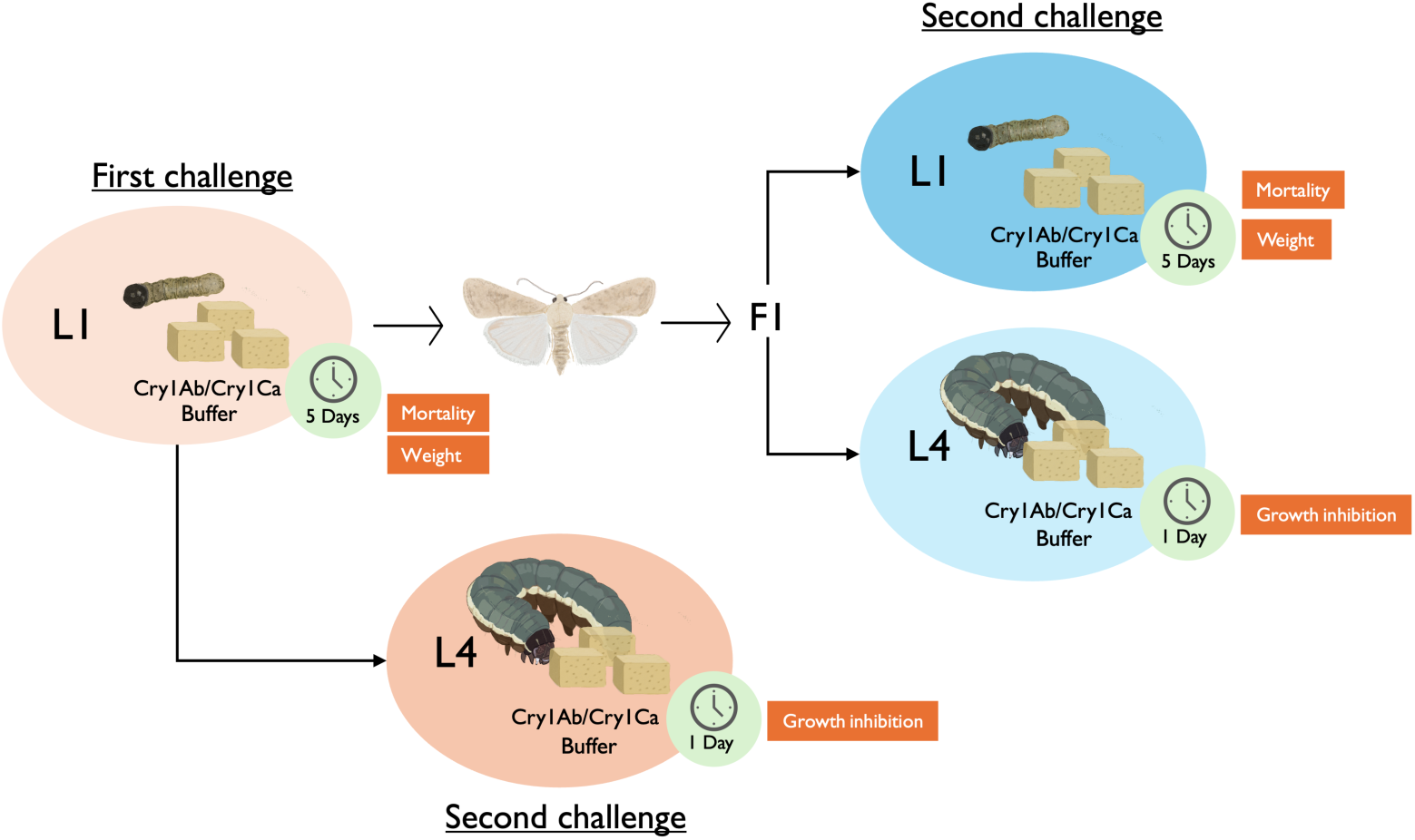
Experimental design.

### 2.5 Statistical analyses

Mortality rates were adjusted using Abbott’s correction formula. Statistical analyses were performed employing a nonparametric methods: the Mann-Whitney U test for two group comparisons and the Kruskal-Wallis test followed by Dunn’s multiple comparisons tests for analyses involving more than two groups. All analyses were carried out using GraphPad Prism version 9.4.1 (GraphPad Inc., USA).

## 3 RESULTS

### 3.1 Sublethal effects on *Spodoptera exigua* L1 instar larvae to Cry1 proteins during first exposure

To observe the sublethal effects on L1 larvae caused by Cry1Ab and Cry1Ca exposure, larvae were exposed for 5 days to different concentrations below the expected LC_50_ for each protein. As anticipated, the mortality levels observed for the L1 larvae exposed to the different concentrations of Cry1Ab or Cry1Ca did not exceed 15% (Figure 2A and 2C). After 5 days of exposure, relative larval weight was used as indicator to sublethal intoxication. Exposure to Cry1Ab resulted in a weight reduction of more than 50%, with a progressive decline in mass as the protein concentration increased (Figure 2B). On the other hand, larvae exposed to Cry1Ca exhibited a moderate weight reduction with approximately a 50% decrease only at the highest concentration used (Figure 2D). Sublethal exposures to Cry1 proteins significantly impacted larval size and weight, resulting in developmental delay. After exposure, treated larvae remained at earlier instars (L1-L2), whereas control larvae had reached the third-instar larvae (L3), an except in the treatment with 1 ng/cm^2^ Cry1Ca, where the larval development was comparable to the control. After transferring the exposed larvae to non-treated diet, the exposed larvae resumed normal behavior (Table 1), indicating that the negative effect caused by Cry1 exposure was not permanent. The mean duration of each life stage, as well as the percentage of individuals that successfully reached the pupal and adult stages, did not differ among treatments and proteins, indicating no fitness costs associated with Cry1 exposure. However, some exceptions were observed, such as larvae treated with 75 ng/cm^2^ of Cry1Ab, where development to pupal stage was delayed. Additionally, exposure to 10 ng/cm^2^ of Cry1Ca reduced the proportion of induvial that successfully pupated.

**Fig. 2.**
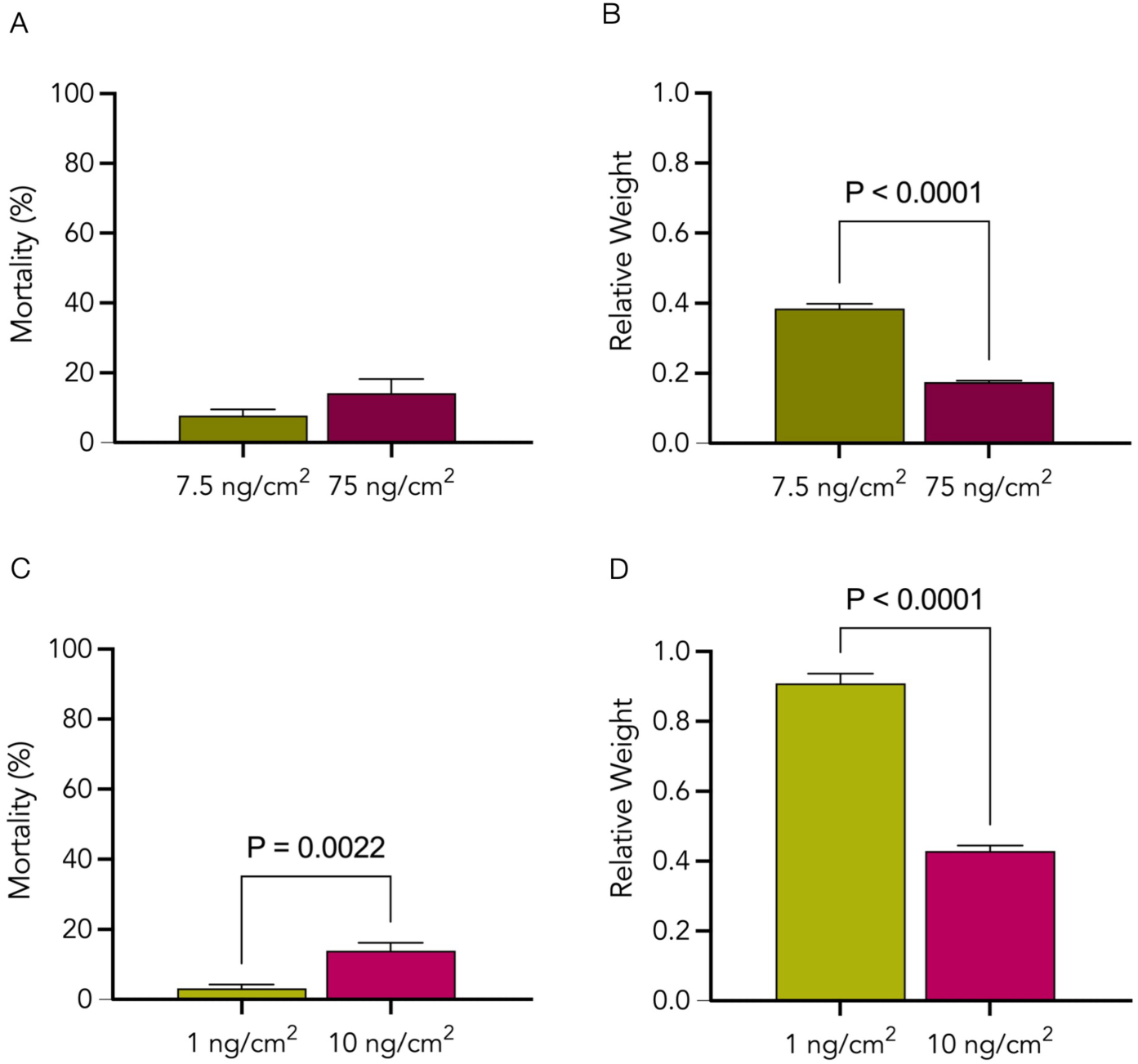
Effect of initial Cry1Ab or Cry1Ca exposure on larval weight and mortality in *Spodoptera exigua*. Larval mortality (A, C) and relative weight (B, D) of *S. exigua* after five days of exposure to different protein concentrations of Cry1Ab (A, B) or Cry1Ca (C, D). Control: L1 larvae exposed to the buffer in which the protein was solubilized.

**Table. 1.** Developmental parameters and survival of *Spodoptera exigua* after first exposure to Cry1Ab or Cry1Ca.

| Parameter | Exposure to Cry1Ab |  |  | Exposure to Cry1Ca |  |
| --- | --- | --- | --- | --- | --- |
|  | Control | 7.5 (ng/cm<sup>2</sup>) | 75 (ng/cm<sup>2</sup>) | 1 (ng/cm<sup>2</sup>) | 10 (ng/cm<sup>2</sup>) |
|  | Mean ± s.e. | Mean ± s.e. | Mean ± s.e. | Mean ± s.e. | Mean ± s.e. |
| **Development time** |  |  |  |  |  |
| Days to pupation from neonate | 13.6 ± 0.5 | 13.7 ± 0.3 | 16.0 ± 0.6* | 15.0 ± 0.6 | 15.7 ± 0.9 |
| Days to adulthood from pupa | 8.1 ± 0.4 | 9.0 ± 1.0 | 7.7 ± 1.2 | 8.7 ± 0.7 | 8.7 ± 0.7 |
| Days to egg-laying after adulthood | 3.1 ± 0.3 | 2.0 ± 0.0 | 2.3 ± 0.8 | 3.0 ± 1.0 | 3.3 ± 0.9 |
| Days to hatching from egg | 2.7 ± 0.2 | 2.7 ± 0.3 | 2.7 ± 0.7 | 2.3 ± 0.3 | 2.7 ± 0.7 |
| **Pupal and adult emergence rates** |  |  |  |  |  |
| Percentage of pupae from larvae | 92.2 ± 1.7 | 85.2 ± 8.5 | 85.1 ± 5.2 | 86.1 ± 1.8 | 71.8 ± 3.8* |
| Percentage of adults from pupae | 88.7 ± 2.4 | 90.6 ± 5.2 | 90.1 ± 6.8 | 92.0 ± 2.6 | 91.7 ± 4.3 |

\* Indicates significant differences (p<0.05) between the control and treatment

### 3.2 Effects of prior exposure to sublethal concentrations of Cry1 proteins on growth inhibition in L4 instar *Spodoptera exigua* larvae after a second exposure

As expected, both Cry1 protein inhibited *S. exigua* larval growth in fourth-instar larvae (L4), in a dose-dependent manner (Figure 3). Specifically, exposure to 7.5 ng/cm² of Cry1Ab resulted in a 52% inhibition, reaching 89% at 750 ng/cm². For Cry1Ca, growth inhibition was 42% at 10 ng/cm² and increased to 95% at 1000 ng/cm². In the case of Cry1Ab (Figure 3A), a comparative analysis between the control group (larvae without prior exposure) and larvae previously exposed as neonates revealed no statistically significant differences, indicating that early-stage exposure did not alter tolerance levels in the same generation.

**Fig. 3.**
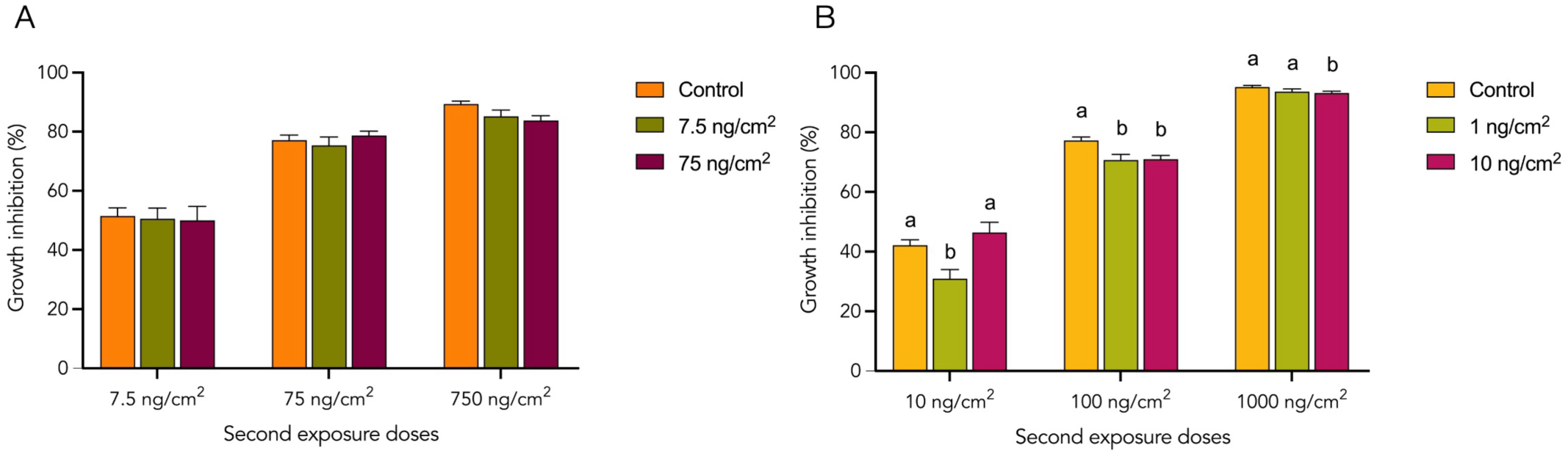
Growth inhibition of the 4^th^ instar *Spodoptera exigua* larvae following second exposure to different concentrations of Cry1Ab (panel A) and Cry1Ca (panel B). Each bar represents the first treatment administered to L1 larvae, while each group corresponds to the concentrations used to exposed L4 larvae for 24 hours, within the same generation. Control: L1 larvae exposed to the buffer in which the protein was solubilized. Lowercase letters indicate significant differences (p<0.05) between the control and different concentrations of the protein.

By contrast, Cry1Ca exhibit a different pattern (Figure 3B). In general trend, larvae treated as neonates showed a significant reduction in growth inhibition compared to the control group. Neonates initially exposed to 1 ng/cm² and then to 10 ng/cm² exhibited a decrease in growth inhibition from 42% to 31% relative to the control and from 77% to 71% when were exposed to 100 ng/cm. Conversely, neonates initially exposed to 10 ng/cm², showed a slightly change in terms of growth inhibition from 77% to 71% at the 100 ng/cm² treatment, and from 95% to 93% at the 1000 ng/cm² treatment.

### 3.3 Changes in the toxicological response of *Spodoptera exigua* larvae over time after sublethal exposure to Cry1 proteins of parentals

To evaluate potential changes in the toxicological response following exposure to Cry proteins, descendant larvae were exposed to the same sublethal concentrations used for the parentals, as well as to an additional, higher concentration. The average weight and mortality in L1 instar larvae were measured after five days of exposure (Figure 4). For Cry1Ab exposure (Figure 4A and 4B), the results revealed no significant differences between control groups and descendants of treated parents across all tested concentrations with both weight and mortality rates remaining consistent. Similar results were observed after exposure of S. *exigua*, descendant L1 larvae to Cry1Ca (Figure 4C and 4D). Interestingly, a reduction in weight, approximately 30% compared to the control group, in larvae descended from parents exposed to 10 ng/cm² was observed. This weight reduction was presented only when larvae were exposed to 10 ng/cm² and not to the other two concentrations tested. No significant differences in terms of mortality were observed between treatments and the control group.

**Fig. 4.**
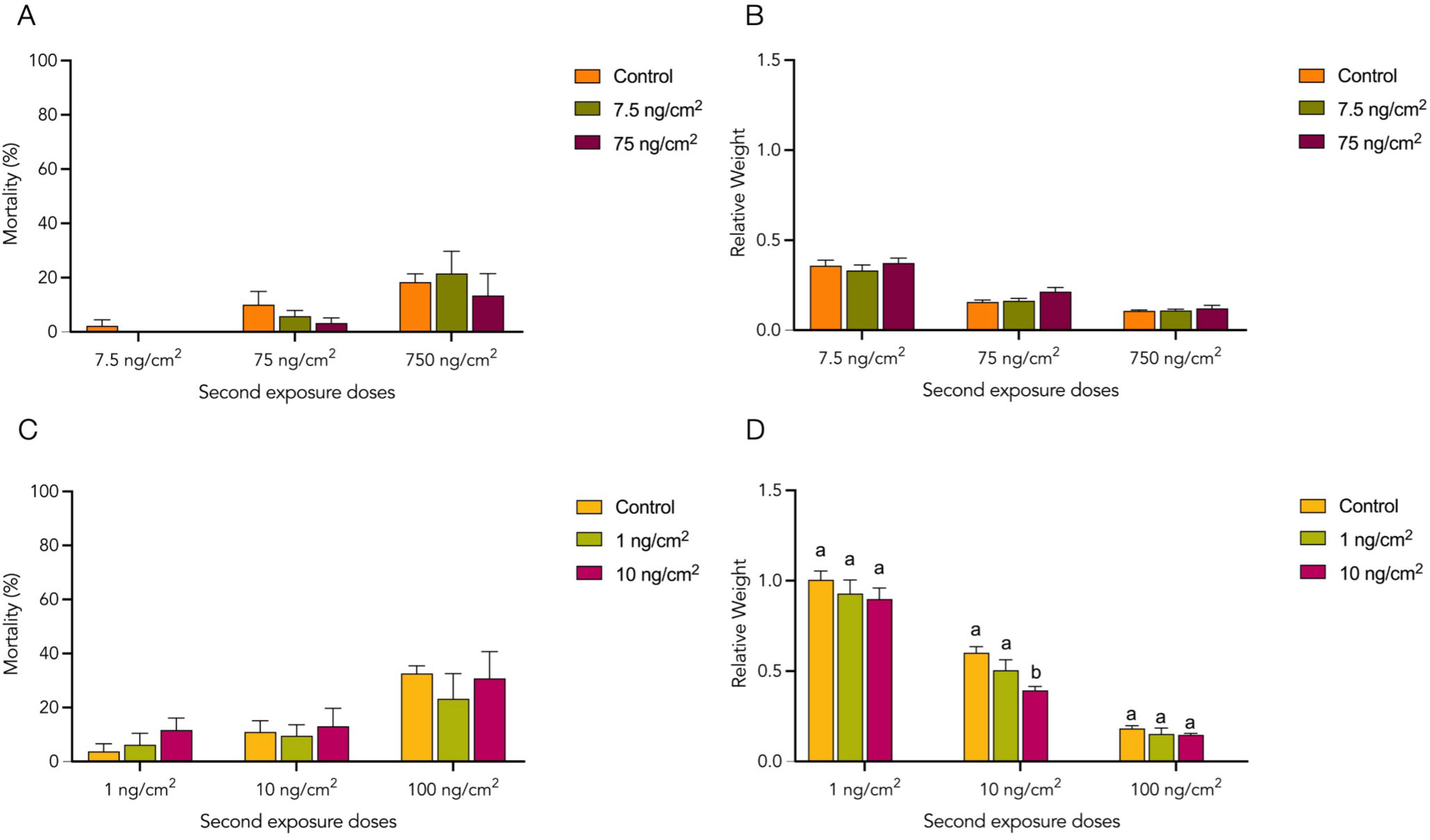
Effect of second Cry1Ab (A, B) or Cry1Ca (C, D) exposure on larval weight and mortality in descendants of *Spodoptera exigua*. Larval mortality (A,C) and relative weight (B,D) of L1 descendants after exposure. Each bar represents the treatment administered to the parental L1 larvae, while each group corresponds to concentration used to expose the descendant L1 larvae for 5 days. Control: L1 larvae exposed to the buffer in which the protein was solubilized. Lowercase letters indicate significant differences (p<0.05) between the control and different concentrations of the protein.

For the assays carried out using descendant larvae at L4 instar, larvae derived from parents exposed to higher concentrations of Cry1Ab exhibited different responses (Figure 5A). When the offspring were exposed to a low dose of 7.5 ng/cm^2^, growth inhibition after 24 hours was reduced, decreasing from 45% to 34% compared to the control. In contrast, when larvae were exposed to the highest dose of 750 ng/cm^2^ the effect was reversed, with growth inhibition increasing from 88% to 93%. These results suggest the existence of a threshold at which some benefit can be observed, further increases in protein exposure led to detrimental outcomes, rather than improvement. When the descendant L4 larvae were treated with Cry1Ca for 24 hours, growth inhibition was similar to the control group (Figure 5B).

**Fig. 5.**
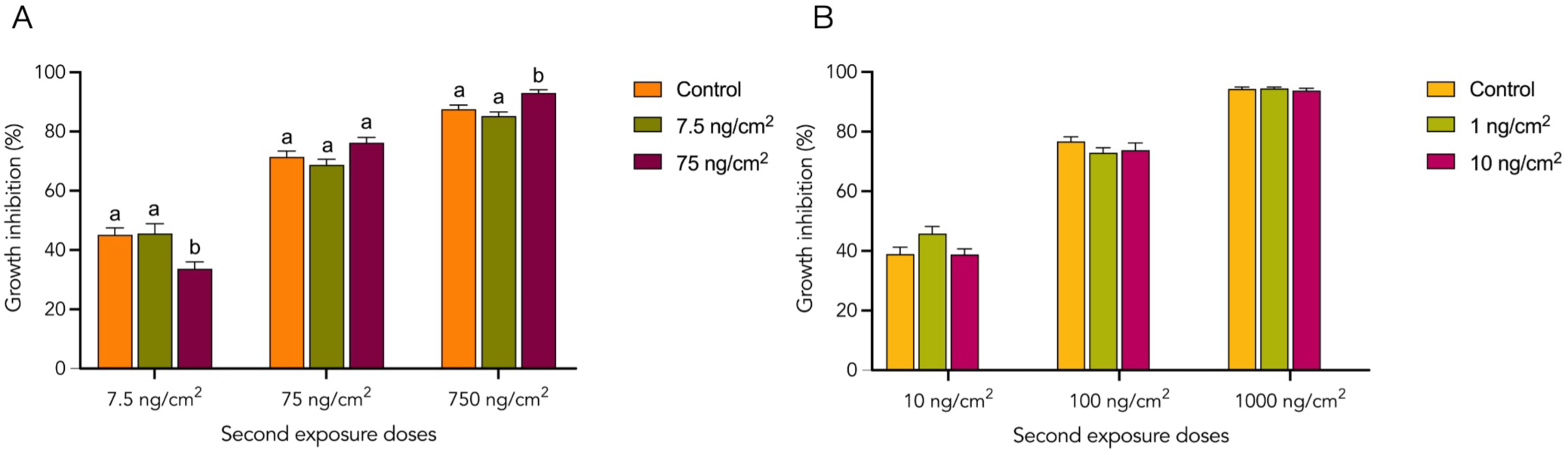
Growth inhibition of the 4^th^ instar *Spodoptera exigua* larvae descendants following parental exposure to different concentrations of Cry1Ab (panel A) and Cry1Ca (panel B). Each bar represents the treatment administered to the parental L1 larvae, while each group corresponds to the concentrations used to expose the descendant L4 larvae for 24 hours. Control: L1 larvae exposed to the buffer in which the protein was solubilized. Lowercase letters indicate significant differences (p<0.05) between the control and different concentrations

## 4 DISCUSSION

The use of Bt-derived products, whether through formulations or transgenic plants, represents a significant advance in pest control technology. However, inappropriate use of these tools threatens their long-term efficacy and sustainability. One of the risks is the potential reduction in insect susceptibility to Bt products. Numerous studies have demonstrated that repeated exposure to Bt vegetative cells, spores and crystals or heat-killed bacteria preparations can enable insects to develop mechanisms that enhance their tolerance.^33–39^ However, little is known about this phenomenon in *S. exigua*, a key agricultural pest. In this context, our study aimed to explore the impact of repeated oral exposure to purified Bt Cry1 proteins in *S. exigua*, an unexplored approach given the scarcity of studies isolating the impact of Cry proteins from other Bt components. To closely simulate field conditions, we used an oral challenge model, in contrast to the more commonly used direct hemocoel injection method. This approach mimics natural exposure scenarios, where the protein needs to overcome various barriers and insect defense mechanisms to reach its target site.^30^

Using this exposure model, we investigated whether there was an increase in tolerance within the same generation after repeated challenges. Rather than using mortality assays, we employed growth inhibition assays, as the larvae were in their fourth instar at the time of the second exposure. At this developmental stage, growth inhibition is a more reliable indicator of toxin impact than mortality, as older instars are less susceptible, requiring notably higher toxin concentrations to induce lethal effects. Growth inhibition, therefore, serves as an alternative measure of toxin susceptibility and has been used in previous studies.^54–57^

Our findings reveal that prior exposure to Cry1Ab did not result in significant changes in tolerance to subsequent Cry1Ab challenges. This suggests that *S. exigua* does not develop an increased tolerance to Cry1Ab within the same generation. In contrast, exposure to Cry1Ca led to a significant decrease in growth inhibition, indicating an enhanced tolerance to this protein. However, this response was not consistent across all tested concentrations, suggesting dose-dependent effect. Similar patterns have been reported in other studies, where minimal changes occur at low initial doses, while higher doses can overwhelm the system, causing physiological damage and impairing the insect’s defense against subsequent exposures. ^40,58^

The differing responses to the two Cry1 proteins could be related to their distinct effects on *S. exigua*. Specifically, Cry1Ab shows moderate insecticidal activity, while Cry1Ca exhibits more potent toxicity, which can lead to higher mortality rates.^55^ Furthermore, previous studies have shown that sublethal exposure to Cry1Ca induces significant physiological changes, including alterations in immune gene expression.^48^ These changes may induce short-term adaptive responses in the insect that potentially contribute to the increased tolerance observed upon subsequent exposure. However, this response may not be solely due to immune changes; other physiological or environmental factors could also be involved.^59^

It has been shown that *S. exigua* can improve survival against several pathogens after prior exposure, via hemocoelic injection, to bacterial components. However, unlike the findings of Haraji et al.,^47^ in this case, prior exposure to isolated proteins only confers a temporary advantage in specific scenarios. The nature of the exposure agent, whether a mixture of molecules or a single component, and the inoculation method are crucial in shaping the response. Similarly, Länger et al.^60^ demonstrated that exposure to the Cry3Aa toxin could induce increased tolerance in *T. molitor* but observed that this response was less pronounced when compared to its effect in combination with other molecules. The use of whole bacterial cells or complete pathogen formulations may then provide greater protection, as insects have evolved over time to detect and respond to such conserved elements. In general, the insect’s immune system can recognize bacterial pathogens, often by surface protein recognition. In *G. mellonella*, direct hemolymph injection of the purified protein LPS induces protective responses against lethal doses of *Photorhabdus luminescens* TT01. However, the injection of PirA_2_B_2_ toxin did not enhance resistance to lethal bacterial infection.^61^ Our findings support the idea that exposure to isolated proteins alone is insufficient to elicit such protective responses; instead, the insect probably requires additional molecular signals to recognize the pathogen and mount a priming defense.

When examining transgenerational effects, we observe a global scenario in which previous exposures generally doesn’t have an impact on offspring, with some exceptions. In contrast to what has been observed in other species, where parental exposure to Bt resulted in increased survival or tolerance in the offspring,^41–43,45^ the tolerance induced by Cry1Ca in this study was not transmitted to the next generation. A difference in our study is that we exposed larvae at early developmental stages, while many of the previously mentioned studies focused on exposing adult insects. Similar results were found by Schulz et al.,^62^ where oral exposure of *T. castaneum* larvae to Bt did not confer any transgenerational benefits in terms of tolerance, contrasting with the effects of adult injection. This suggest that the developmental stage at which exposure occurs plays a critical role in influencing transgenerational effects.

In addition, the development of tolerance mechanisms can often incur significant fitness costs in the parental generation, which may negatively affect offspring development. For example, in *T. molitor*, offspring exhibited prolonged developmental times or reduced weight at the pupal stage following exposure.^44^ Similarly, studies in *T. castaneum* have reported reduced fecundity and prolonged development.^42,62^ In the case of *S. exigua* exposed to Cry1Ca, while there was some evidence of increased tolerance in the parental generation at certain concentrations, the subsequent generation showed a reduced tolerance at the higher sublethal concentrations. Additionally, parental individuals exposed to these concentrations showed a decrease in pupation rate. For Cry1Ab, parental exposure to higher concentrations resulted in delayed developmental time and greater growth inhibition in the offspring, suggesting that sublethal exposure not only failed to increase tolerance, but may have compromised the fitness and resistance in the next generation. Interestingly, a reduction in growth inhibition was observed when the second exposure involved lower concentration, demonstrating the dose-dependent response involved in this process. These results highlight the complexity of transgenerational tolerance mechanisms. While initial exposures may provide short-term benefits, they may also lead to increased susceptibility in subsequent generations.

The differences observed here were found under highly controlled laboratory conditions, which may not fully capture the complexity of field environments where multiple factors interact, such as plant-insect interactions, exposure duration, toxin concentrations, exposure to different combinations of proteins, degradation factors, etc.^63^ Although statistical differences were found, they may not be biologically significant, as the effects were minimal and the fact that these responses were neither generalized nor sustained over time. Therefore, based on the current data, the risk of reduced efficacy associated with the use of purified Cry proteins for controlling *S. exigua* appears to be minimal.

## 5 CONCLUSION

We orally exposed *S. exigua* to the purified proteins of Bt, Cry1Ab and Cry1Ca, and analyzed changes following a second exposure within the same generation and in subsequent generation. The intragenerational analysis showed that pre-exposure to Cry1Ab did not affect tolerance to subsequent challenges. However, Cry1Ca induced a slight increase in tolerance under certain conditions. Furthermore, transgenerational analysis revealed no significant increase in tolerance in offspring. These results suggest that while initial exposures to sublethal doses of Bt proteins may induce minor physiological changes, such responses are insufficient to drive the development of long-term resistance, particularly in a field-relevant context. Future studies should investigate the interactions between crystals and spore preparations, as these may provide additional stimulation mechanisms. Understanding these interactions will help to refine Bt-based pest management strategies and ensure their long-term future effectiveness.

## ACKNOWLEDGEMENTS

This research was funded by grants from the Spanish Ministry of Science and Innovation PID2021-122914OB-I00 (co-funded by EU FEDER funds) and a PROMETEO 2024 program from the Conselleria de Educación, Cultura, Universidades y Empleo, Generalitat Valenciana, Spain (CIPROM/2023/56). S.V. is a beneficiary of a Generalitat Valenciana GRISOLIA grant (GRISOLIAP/2021/046).

## DATA AVAILABILITY STATEMENT

The data that support the findings of this study are available from the corresponding author upon reasonable request.

## Footnotes

1 This paper was presented at a scientigic meeting

